# CD14 and TLR4 contribute to the circadian regulation of retinal phagocytosis as co-receptors

**DOI:** 10.64898/2026.02.26.708367

**Authors:** Leila Dhaoui Hajem, Julie Enderlin, Quentin Rieu, Sara Krim, Jennyfer C. Parnasse, Clément Materne, Geneviève Marcelin, Thierry Huby, Emeline F. Nandrot

**Author notes:** Corresponding author: Emeline F. Nandrot, Institut de la Vision, 17 rue Moreau, Paris, F-75012, France.

## Abstract

Retinal pigment epithelium (RPE) cells perform crucial functions for vision, among which the daily clearance of photoreceptor outer segment (POS) oxidized extremities. POS phagocytosis is under circadian regulation, peaking only once a day despite the constant contact between both cell types. Alphavbeta5 integrin receptors and MFG-E8 ligands synchronize POS phagocytosis and activate the MerTK internalization receptor via an intracellular signaling cascade. Recently, we identified scavenger receptors CD36 and SR-B2/LIMP2 as POS internalization regulators. We now highlight that innate immunity receptors CD14 and TLR4 interact with POS as stimulatory coreceptors in a tissue-specific fashion. CD14 and TLR4 associate partially with lipid rafts, and their activation triggers MyD88-dependent JNK and ERK1/2 (p44/42) kinases. *In vivo*, CD14 and TLR4 protein levels are replenished in the hours leading to the phagocytic peak. In addition, the phagocytic peak is lost in *Tlr4^-/-^* RPE cells, thus confirming that TLR4 regulates this function. Finally, CD14 and TLR4 associate with SR-B2, partner with CD36 and MerTK, highlighting that several receptors contribute together to the fine regulation of POS phagocytosis as a macromolecular machinery.

## Introduction

Clearance of apoptotic cells is essential for tissue homeostasis. In the body, two categories of phagocytes are required for this task: specialized phagocytes such as macrophages, and non-specialized phagocytes, such as Sertoli and retinal pigment epithelium (RPE) cells (Finnemann and Rodriguez-Boulan, 1999). The RPE is a monolayer of hexagonal polarized post-mitotic cells. It performs functions crucial for photoreceptor survival and retinal homeostasis, among which the phagocytosis of aged photoreceptor outer segments (POS) (Strauss, 2005; Lakkaraju et al, 2020). If POS phagocytosis does not occur it will lead to the development of early onset rod-cone dystrophies in animals and humans (Bok and Hall, 1971; Gal et al, 2000; Nandrot et al, 2000; Parinot and Nandrot, 2016; Audo et al, 2018), while if phagocytosist is dysregulated it will lead to late onset pathologies such as age-related macular degeneration (AMD) (Nandrot et al, 2004).

The molecular machinery tackled during POS phagocytosis is similar to the one used by macrophages to clear apoptotic cells, based on the recognition by receptors at RPE apical cell surface of an “eat-me” signal –phosphatidylserines (PtdSer)– exposed on the outer leaflet of POS membranes (Ravichandran, 2011; Ruggiero et al, 2012). PtdSer are first recognized by αvβ5 integrin receptors (Finnemann et al, 1997; Nandrot et al, 2004), then POS extremities are internalized by MerTK receptors (D’Cruz et al, 2000; Nandrot et al, 2000). MerTK receptors also control the amount of POS that can be attached to αvβ5 integrin dimers, likely via the cleavage of its extracellular domain that appear to be essential to its regulation, and thus to the phagocytic process, both *in vitro* and *in vivo* (Nandrot et al, 2012; Law et al, 2015; Enderlin et al, 2024). A third family of receptors is implicated in this machinery, scavenger receptors such as CD36 and SR-B2/LIMP2, that have been shown to regulate the speed of POS internalization (Finnemann and Silverstein, 2001; Rieu et al, 2022). PtdSer are recognized either directly by receptors expressed at the RPE surface, e.g. scavenger receptors (Ryeom et al, 1996a; Ryeom et al, 1996b; Rieu et al, 2022), or indirectly by ligands expressed in the interphotoreceptor matrix serving as bridge molecules, notably MFG-E8 and Gas6/Protein S, respective ligands of αvβ5 integrin and MerTK receptors (Hall et al, 2005; Nandrot et al, 2007; Parinot et al, 2024).

For RPE cells the “eat-me” signal is not sufficient to phagocytose POS (Lew et al, 2014). Despite the constant contact between RPE cells and photoreceptors, POS phagocytosis is rhythmic and follows a circadian rhythm. In fact, POS phagocytosis occurs once a day and the peak is registered 1.5-2hrs after light onset for rod photoreceptors (LaVail, 1976; Nandrot et al, 2004). The whole machinery is synchronized by the αvβ5/MFG-E8 couple which subsequently activates the MerTK receptor (Nandrot et al, 2004). Lately, we demonstrated yet another difference between RPE cells and macrophages in the opposite roles of Gas6 and Protein S on the regulation of POS phagocytosis and sMerTK release (Nandrot, 2018; Parinot et al, 2024).

Unlike macrophages that circulate and eliminate apoptotic cells when they encounter them, RPE cells and POS are constantly closely interacting. However, phagocytosis occurs only once a day thus suggesting a tight molecular control of the RPE phagocytic machinery to a stronger extent than for macrophages to limit phagocytosis outside of peak activity time. Interestingly, RPE cells express a battery of receptors at their apical surface that have been studied on macrophages but their participation in POS phagocytosis has not been investigated yet. Among those figure the Cluster of Differentiation 14 (CD14) and Toll-Like Receptor 4 (TLR4) (Sun et al, 2006; Finnemann et al, 1997).

CD14 is a 55kDa glycosyl phosphatidylinositol (GPI)-anchored glycoprotein existing under two forms: a membrane-bound form (mCD14) and a soluble form (sCD14) (Wu et al, 2019). The membrane bound form is anchored to the cell membrane via a GPI tail. The soluble form is secreted by activated cells and is released by proteinase-dependent (48kDa) and - independent shedding (55kDa). Monocytes, macrophages, dendritic cells and microglia, as well as hepatocytes, fibroblasts, epithelial and endothelial cells express mCD14 (Wu et al, 2019). In addition, Elner and colleagues have shown that CD14 is expressed by cultured human primary RPE cells (Elner et al, 2003). CD14 can be activated by LPS (Wright, 1999). CD14 is implicated is the clearance of apoptotic cells by macrophages, and its absence in *CD14^-/-^* mice leads to a defect in the elimination of apoptotic cells (Gregory and Devitt, 1999; Devitt et al, 2003; Devitt et al, 2004).

TLR4 is a pattern recognition receptor expressed by immune cells such as dendritic cells and macrophages, and non-immune cells like epithelial cells (Kawazaki and Kawai, 2014; Ciesielska et al, 2021). TLR4 is implicated in innate immunity as proinflammatory reaction activator since its activation leads to cytokine and chemokine factors transduction (Matveichuk et al, 2024). TLR4 expression by RPE cells has been confirmed, and it has been shown to play an important role in the inflammatory response in AMD (Klettner and Roider, 2021).

CD14 also functions as a co-receptor for TLR4 in the pro-inflammatory immune response (Elner et al, 2003; Ciesielska et al, 2021). CD14 and TLR4 are considered as the primary receptors for bacterial lipopolysaccharides (LPS) (Ciesielska et al, 2021; Chow et al, 1999). The activation of TLR4 starts when CD14 interacts with LPS particles via the liposaccharide binding protein (LBP) (Kim et al, 2005; Wu et al, 2019; Ciesielska et al, 2021). Once fixed on LPS, CD14 interacts with the MD2 protein to activate TLR4. In contrast to other TLRs, TLR4 activation triggers two different signaling pathways resulting in a pro-inflammatory response via the secretion of cytokines (TNFα, IL8, IL6…), chemokines and adhesion molecules (Borzęcka et al, 2013; Ciesielska et al, 2021). These two signalling pathways are dependent on either MyD88 or TRIF (Klettner and Roider, 2021). Interestingly, it has been revealed that some raft proteins, known to potentiate signaling pathways activation, contribute to the activation of the CD14–TLR4 duet (Płóciennikowska et al, 2015). In addition, TLR4 has been previously shown to interact with the CD36 scavenger receptor during POS phagocytosis, thus suggesting other receptors could contribute to a larger macromolecular complex (Finnemann and Silverstein, 2001).

Hence, in this article we set out to characterize the implication of CD14 and TLR4 as well as associated signaling pathways in the regulation of POS circadian phagocytosis *in vitro* and *in vivo*, alone or as coreceptors.

## Results

### CD14 and TLR4 are expressed by RPE cells and associate with POS

In order to characterize their implication in POS phagocytosis, we first wanted to confirm CD14 and TLR4 expression and sizes *in vitro* in J774 macrophages, the rat RPE-J and human ARPE-19 cell lines, as well as *in vivo* in full eyecups, RPE/choroid and retinal tissues (Fig. 1A). Both receptors are expressed by the different cellular cell types *in vitro*. In retinal tissues, CD14 showed a wide expression pattern being expressed in full eyecups, RPE/choroid and retina fractions. Interestingly, in the retina two bands can be observed, most probably rather due to different glycosylation patterns than to clivage given the size observed (close to 48kDa). TLR4 shows a different profile of cellular and tissular expression: while its expression is limited in J774 macrophages and human ARPE-19 cells, it is much more expressed in RPE-J cells. *In vivo*, the receptor expression detected in eyecups seems to be more related to a RPE/choroid than to a retinal expression. As well, different sizes of bands can be observed, potentially linked to both glycosylation and clivage (Yang et al, 2018).

**Figure 1.**
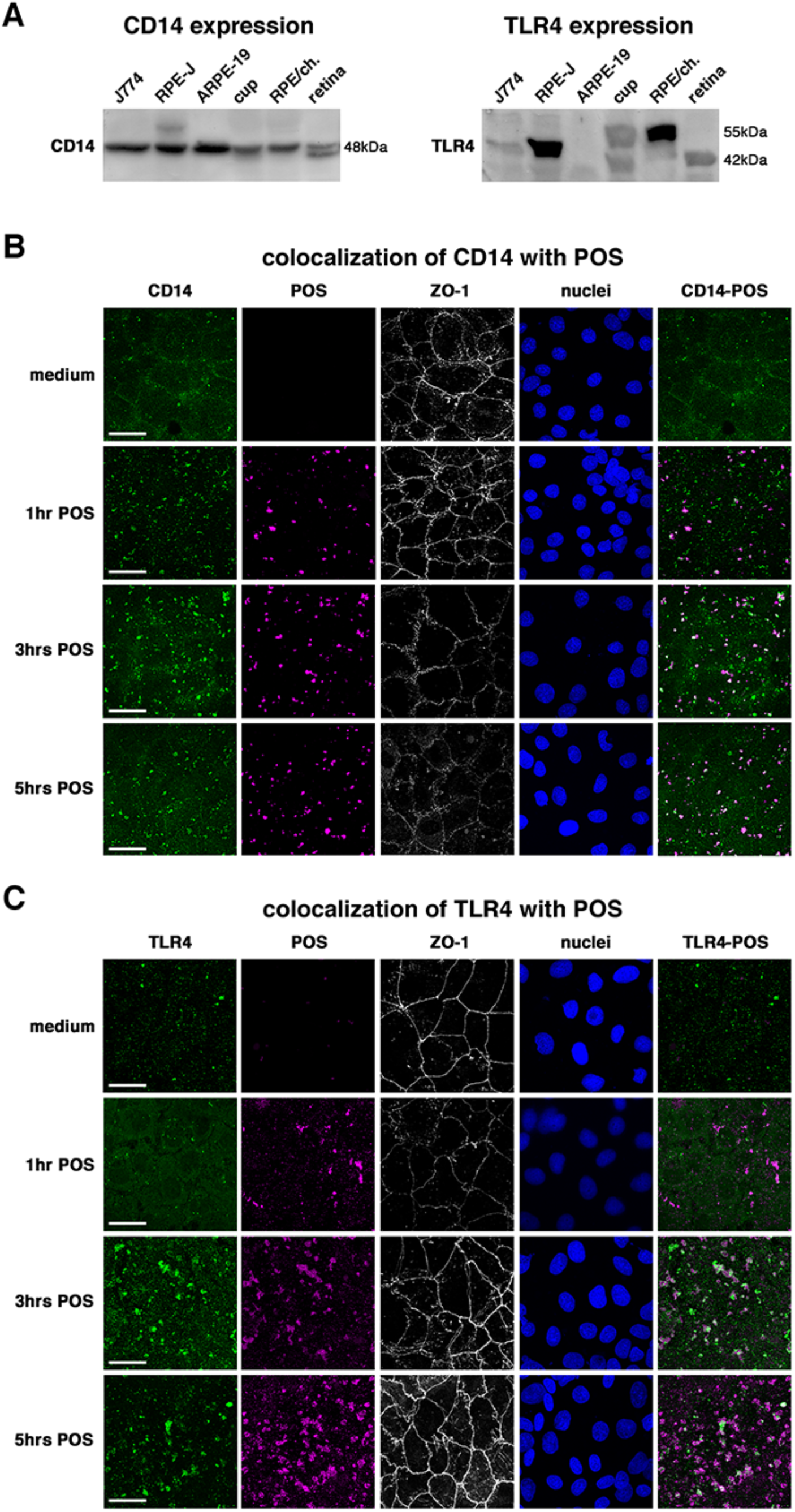
CD14 and TLR expression and colocalization with POS during phagocytosis. **A** Immunoblot expression profiles of CD14 (left) and TLR4 (right) in macrophages (J774), rat (RPE-J) and human (ARPE-19) RPE cells, full eyecups (cup), RPE/choroid (RPE/ch.) and retinal (retina) fractions as indicated. Sizes are indicated in kDa. **B,C** Immunocytochemistry labelings of CD14 (**B**, green) and TLR4 (**C**, green), POS (purple), ZO-1 (grey), nuclei (blue) and receptor–POS overlay after incubation in medium or with POS for 1, 3, and 5 hrs as indicated. Scale bars: 20µm.

Next, we assessed if CD14 and TLR4 recognize and associate with POS during phagocytosis by performing colabeling immunocytochemistry assays at 1, 3 and 5 hours of POS challenge. In normal conditions, i.e. before being challenged with POS, RPE-J cells show a punctate expression pattern of CD14 at the cell surface and at cell junctions (Fig. 1B). Upon POS challenge, the intensity of visible surface CD14 receptors increases at 1 hr and 3 hrs, before decreasing again at 5 hrs. In addition, receptors relocalize in the membrane leaflet to colocalize with POS. This colocalization takes place at all times (1, 3 and 5 hrs of incubation), and reaches its maximum at 3 and 5 hrs.

TLR4 is also organized in marked puncta at the cell surface in medium conditions (Fig. 1C). Similarly to CD14, when cells are fed with POS TLR4 receptors relocate at the cell surface to colocalize with POS, with the biggest association after 3 hrs of POS challenge.

Interestingly, both receptors appear to be organized in clusters during phagocytosis, and the peak of clustering is registered mostly around 3 hrs of POS incubation.

### CD14 and TLR4 associate in RPE cells during POS challenge

To test if both receptors associate during POS phagocytosis as it has been shown in macrophages (Elner et al, 2003; Ciesielska et al, 2021), we repeated the same immunocytochemistry experiments and labeled each receptor simultaneously at different times of POS challenge (Fig. 2A). While some CD14 and TLR4 receptors can be associated in medium conditions, it is not the case for the majority of receptors. However, when stimulated by POS, the receptors association seems to increase rapidly at 1 hr and reaches a peak at 3 hrs, before slowly decreasing 5 hrs.

**Figure 2.**
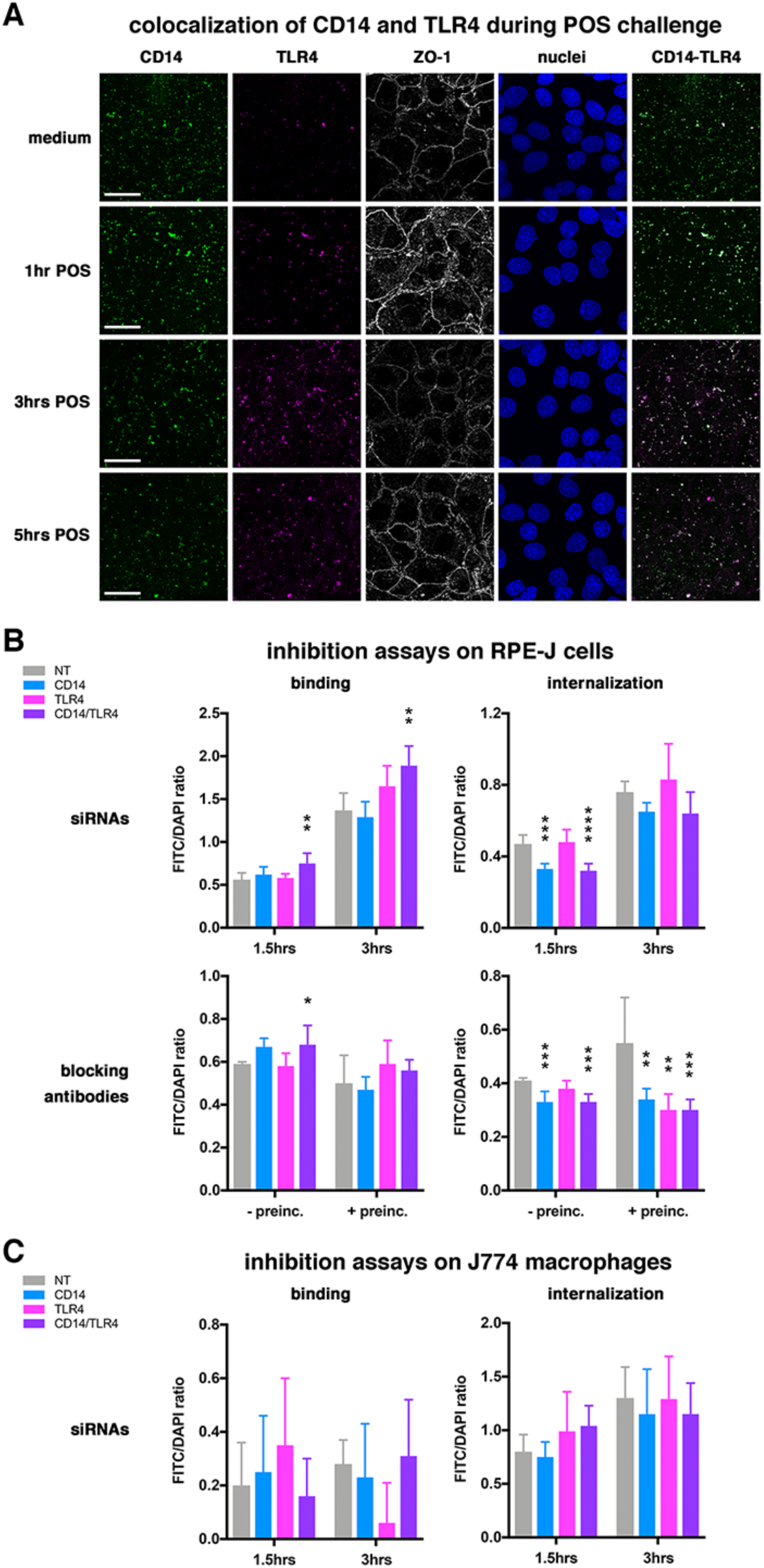
CD14–TLR colocalisation and impact of inhibition assays during POS phagocytosis. **A** Immunocytochemistry labelings of CD14 (green) and TLR4 (purple), ZO-1 (grey), nuclei (blue) and CD14–TLR overlay after incubation in medium or with POS for 1, 3, and 5 hrs as indicated. Scale bars: 20 µm. **B,C** Inhibition assays on RPE-J (**B**) and J774 macrophages (**C**) using siRNAs (**B,C**, 1.5 and 3 hrs) or blocking antibodies (**B**, ± preincubation, 3 hrs), either alone or combined as indicated. CD14 inhibition is in blue, TLR4 inhibition in pink and double inhinbitions assays in purple. N = 4-5 (RPE-J siRNAs), 4-7 (blocking antibodies) and 3-5 (J774 siRNAs). Data information: In (B–C), data are presented as mean ± s.d., * P < 0.05, ** P < 0.01, *** P < 0.001 and **** P < 0.0001 (One-way ANOVA with Dunnett post correction).

### CD14 inhibition directly impacts RPE cells phagocytosis

To assess their exact contribution to RPE cells phagocytosis, we tested whether the inhibition of these receptors had an affect of POS engulment. For that purpose, we performed two kinds of inhibition assays, downregulation of gene expression using siRNAs and blocking receptors function directly at the functional level using antibody inhibition. siRNA inhibition showed different effects for *Cd14* and *Tlr4* (Fig. 2B). *Cd14* inhibition alone decreased significantly POS internalization at 1.5 hrs incubation with a similar trend observed at 3 hrs. However *Tlr4* gene expression did not show any effect on any phagocytic step or time of incubation. When both receptors were blocked at the same time, we observed an increase in POS binding at 1.5 and 3hrs, while the effect on internalization was the same as for the downregulation of *Cd14* alone. The use of blocking antibodies with or without preincubation allows us to see if receptor interaction/association at RPE cell surface are required before the recognition of the target (Finnemann and Silverstein, 2001). Without preincubation, antibody inhibition showed a similar trend at 3 hrs of POS challenge to the one observed with siRNAs: binding was increased when both receptors were blocked, and internalization was diminished when CD14 alone or both receptors were blocked. However, POS internalization was also affected when cells were preincubated with TLR4 antibodies alone.

In order to compare the effect between RPE cells and macrophages, we proceeded with siRNA inhibition assays on J774 macrophages. For these cells, no significant effect was observed at any step or time-point of phagocytosis tested (Fig. 2C).

### CD14 and TLR4 are partially located in lipid rafts

In innate immunity, it is known that the CD14–TLR4 interaction takes place in lipid raft domains of the cell membrane, and that it is promoted by some lipid rafts proteins (Pleiffer et al, 2001; Ciesielska et al, 2021). Our immunocytochemistry data showed that both CD14 and TLR4 organize in clusters during phagocytosis, and the clustering increases with incubation time to reach a peak at 3 hrs of POS challenge (Fig. 1B,C). Lipid rafts being cell signaling platforms, we wanted to confirm whether our receptors colocalize with raft proteins. We reapeated our immunocytochemistry assays using caveolin as lipid raft marker. Association of CD14 with lipid rafts was barely detected at 1 hr of phagocytosis when CD14 receptors already associate with POS (Fig. 3A). The largest association of CD14 with caveolin was observed at 3 hrs, and somewhat at 5 hrs. Interestingly, TLR4 association with lipid rafts was detected as early as 1 hr of POS challenge, and was maintained during the whole time of the experiments (Fig. 3B). For both receptors, we noticed that they only partially colocalize with the raft marker caveolin.

**Figure 3.**
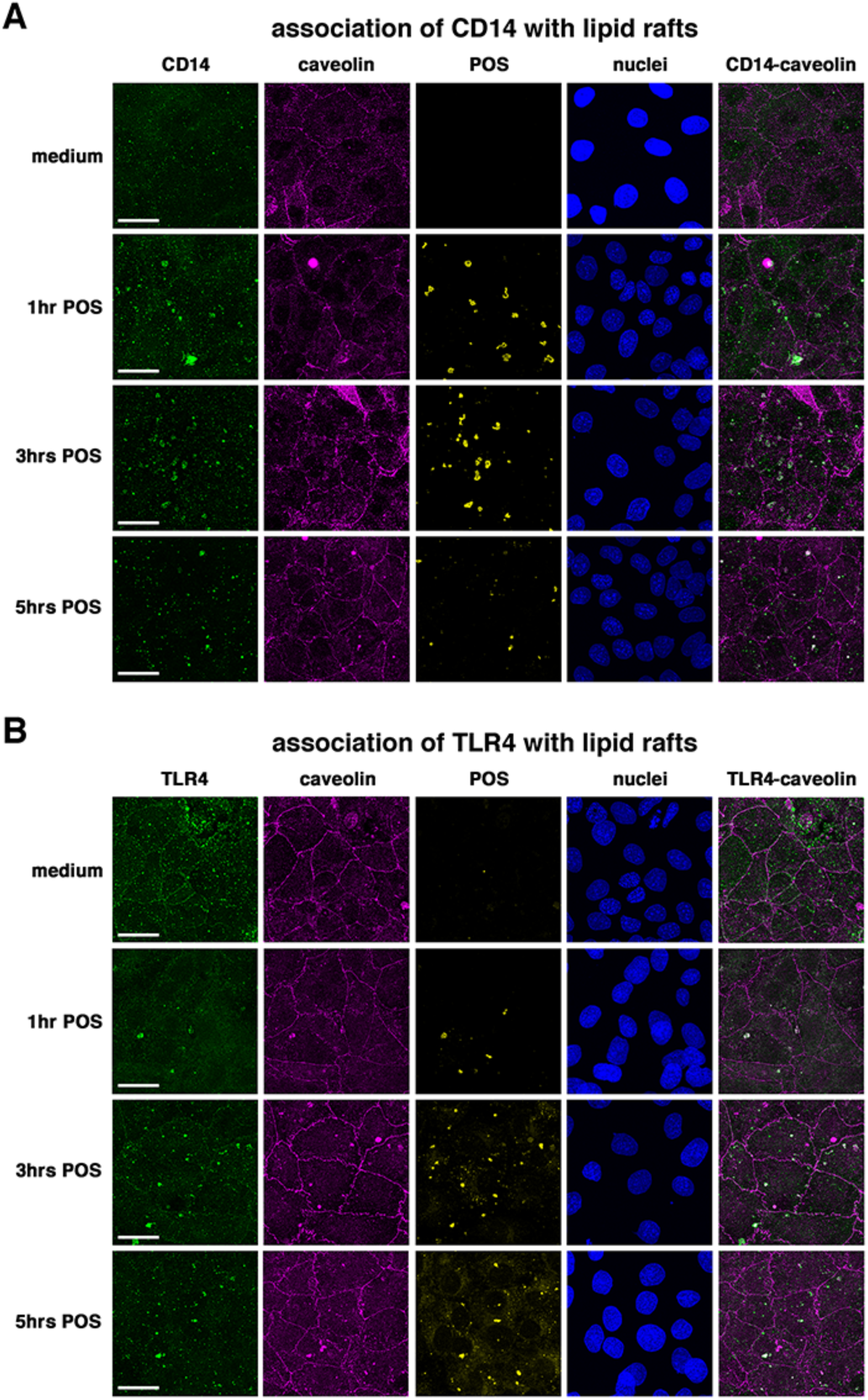
Association of CD14 and TLR4 with lipid rafts. **A,B** Immunocytochemistry labelings of CD14 (**A**, green) and TLR4 (**B**, green), caveolin (purple), POS (yellow), nuclei (blue) and receptor–caveolin overlay after incubation in medium or with POS for 1, 3, and 5 hrs as indicated. Scale bars: 20 µm.

### CD14–TLR4 triggering activates the JNK and ERK1/2 (p44/42) MAPK pathways

We then wanted to characterize which signaling pathway/s is/are activated downstream of this interaction. For that purpose, we repeated our phagocytic challenge experiments at 1, 3 and 5 hrs and tested the expression and phosphorylation of known transduction pathway proteins on immunoblots. CD14 and TLR4 receptors have been shown to be associated with both MyD88-dependent and independent pathways (Ciesielska et al, 2021). The first components of the MyD88-dependent pathway, MyD88, IRAK1 and TRAF6, did not show any difference in their expression profiles in medium conditions or when POS were added (Fig. 4). After quantification, MyD88 show a slighty significant decrease in its whole protein expression levels after 5 hrs of POS challenge. TAK1 is the first phsphorylated protein in this pathway, and it did not show any difference in any condition. Then, this pathway divides into three branches of MAPK proteins, p38, JNK and ERK1/2 (p44/42), these being activated by phosphorylation. No differences were detected in the overall expression of these various proteins. While p38 had an uneven pattern of activation –sometimes increased sometimes decreased–, JNK was strongly stimulated by the addition of POS after 1 hr and then its activation decreased, and ERK1/2 (p44/42) showed a robust increase of its phosphorylation pattern with time of presence of POS.

**Figure 4.**
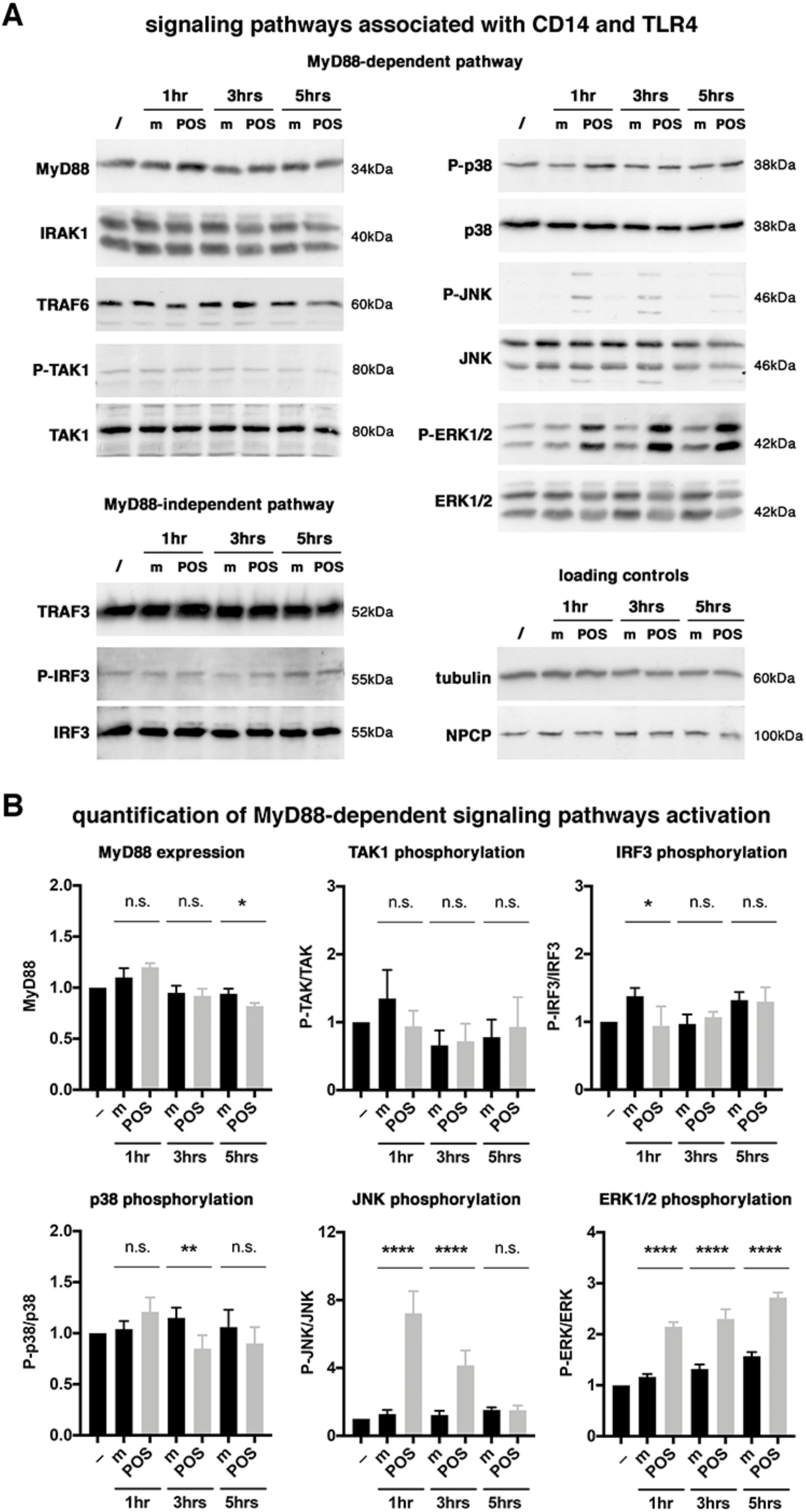
Downstream signaling pathways associated with CD14–TLR. **A** Representative immunoblots of MyD88-dependent (top panels) and MyD88-independent (bottom left panels) signaling pathway members, and loading controls (bottom right panels) expression and phosphorylation (P-) profiles after incubation in medium or with POS for 1, 3, and 5 hrs as indicated. Sizes are indicated in kDa. **B** Bar graphs of corresponding quantifications as indicated are presented as means ± s.d., N = 3-6, n.s. non significant, * P < 0.05, ** P < 0.01, **** P < 0.0001 (One-way ANOVA with Dunnett post correction).

### CD14 and TLR4 are part of a macromolecular machinery

Since interaction of CD14 and TLR4 with other receptors have been identified in innate immunity (Bowdish et al, 2009; Amiel et al, 2011; Taban et al, 2022; Zizzo and Cohen, 2018), we wanted to verify if other identified RPE phagocytosis receptors interact with them as well during POS phagocytosis. We first studied possible the association of CD14 and TLR4 with another regulatory receptor, SR-B2 (Nandrot et al, 2000; Rieu et al 2022; Enderlin et al, 2024). Our results show that CD14 and TLR4 associate strongly with SR-B2, especially at 3 hrs of POS challenge (Fig. 5A). We also recorded an association between MerTK and CD14 mostly at 1 hr, which was weaker.

**Figure 5.**
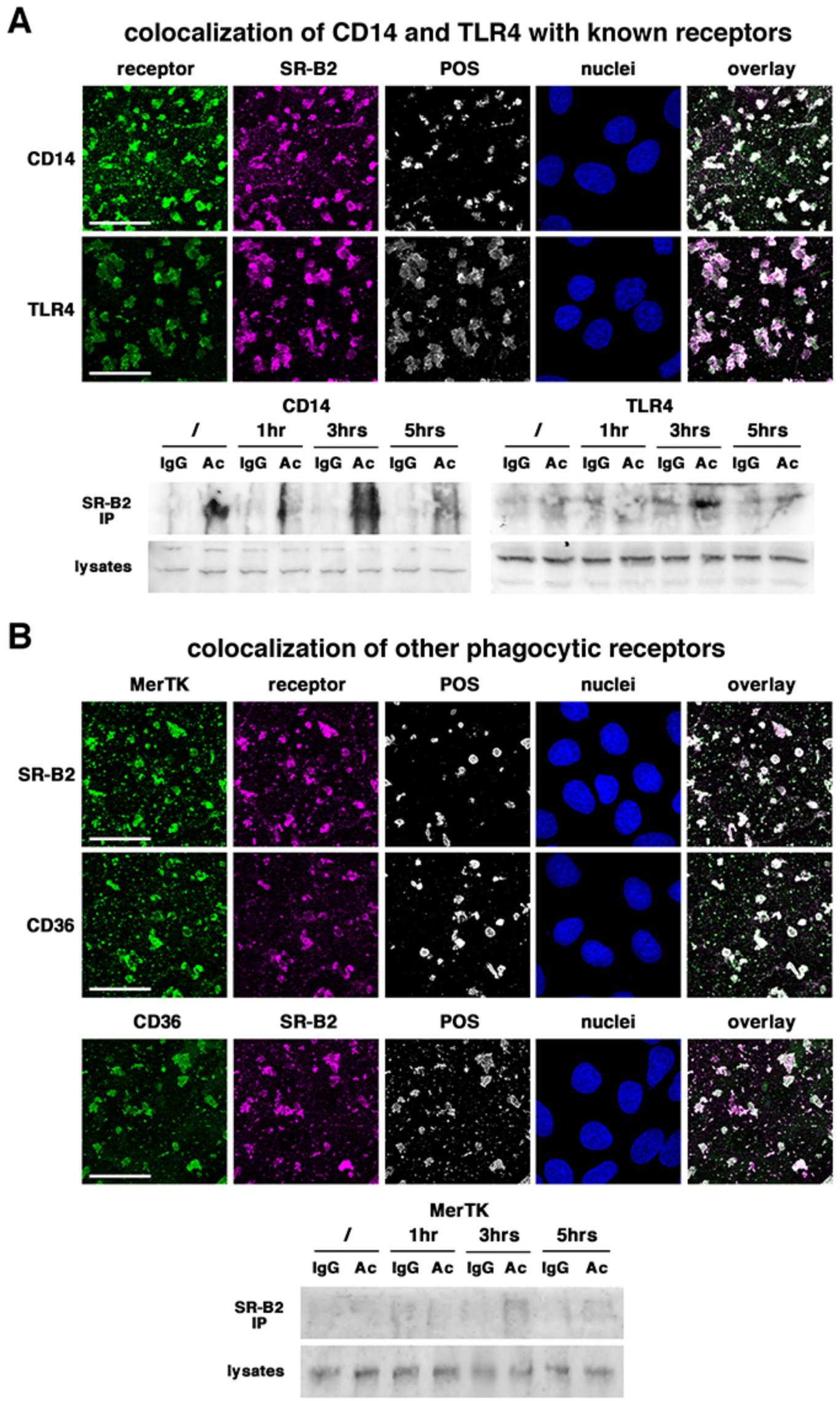
Association with other phagocytic receptors. **A** Immunocytochemistry labelings of CD14 (top, green) and TLR4 (bottom, green) with SR-B2 (red) after 3 hrs of POS (grey) challenge. **B** Immunocytochemistry labelings of MerTK (top and middle, green) with SR-B2 (top, red) and CD36 (middle, red), and of CD14 (bottom, green) with SR-B2 (bottom, red) after 3 hrs of POS (grey) challenge. Nuclei are in blue, scale bars: 20 µm. **A,B** Immunoblots for various receptors of immunoprecipitation assays (top) using non-immune IgG (IgG) or an anti–SR-B2 antibody (Ac) and corresponding lysates (bottom) without (/) or after 1, 3 and 5 hrs incubation with POS, as indicated.

To investigate the regulation of this machinery and further, we assessed the possible interaction of the internalization receptor MerTK with scavenger receptors SR-B2 and CD36 (Finnemann and Rodriguez-Boulan, 1999; Rieu et al, 2022). Interestingly, both receptors associate with MerTK, with a maximum reached after 3 hrs of incubation with POS (Fig. 5B). Concomitantly, CD36 and SR-B2 interact with each other as well.

We performed immunoprecipitation assays to confirm these data using SR-B2 as bait, as it appears to be an important member of this machinery. SR-B2 receptor associate with CD14, TLR4 and MerTK at the different incubation times tested (1, 3 and 5hrs) with a peak registered at 3hrs, while cell lysates do not show any modification in any receptor expression levels (Fig. 5A,B).

### CD14 and TLR4 expression along the light:dark cycle and impact on in vivo phagocytosis

*In vivo*, POS phagocytosis is rhythmic and peaks 1.5-2 hrs after light onset (LaVail, 1976; Nandrot et al, 2004). We thus asked the question if our receptors display a rhythmic expression pattern since other phagocytosis receptors do, such as *MerTK* and scavenger receptors *Marco* and *Cd36* (Law et al, 2015; Rieu et al, 2022). We studied the circadian expression of CD14 and TLR4 on WT and *β5^-/-^* mouse RPE/choroid tissues, *β5^-/-^* mice being considered as acontrol as they present an arrhythmic pattern of phagocytosis (Nandrot et al, 2004). The RNA expression pattern is different between *Cd14* and *Tlr4* (Fig. 6A,B). *Cd14* presents a peak between 4 and 6 hrs, before light onset, and two peaks at 9-11 hrs and at 22 hrs (Fig. 6A). In *β5^-/-^* RPE/choroids, the profile is overall increased, with an exacerbated peak observed at 4-6 hrs and an extra peak at 24 hrs. At the protein level, the CD14 expression profile roughly matches the RNA profile, except at 22 hrs for WT samples when a protein expression increase is not observed and at 4-6 hrs in *β5^-/-^* samples when the expression is not augmented as much as for RNAs.

**Figure 6.**
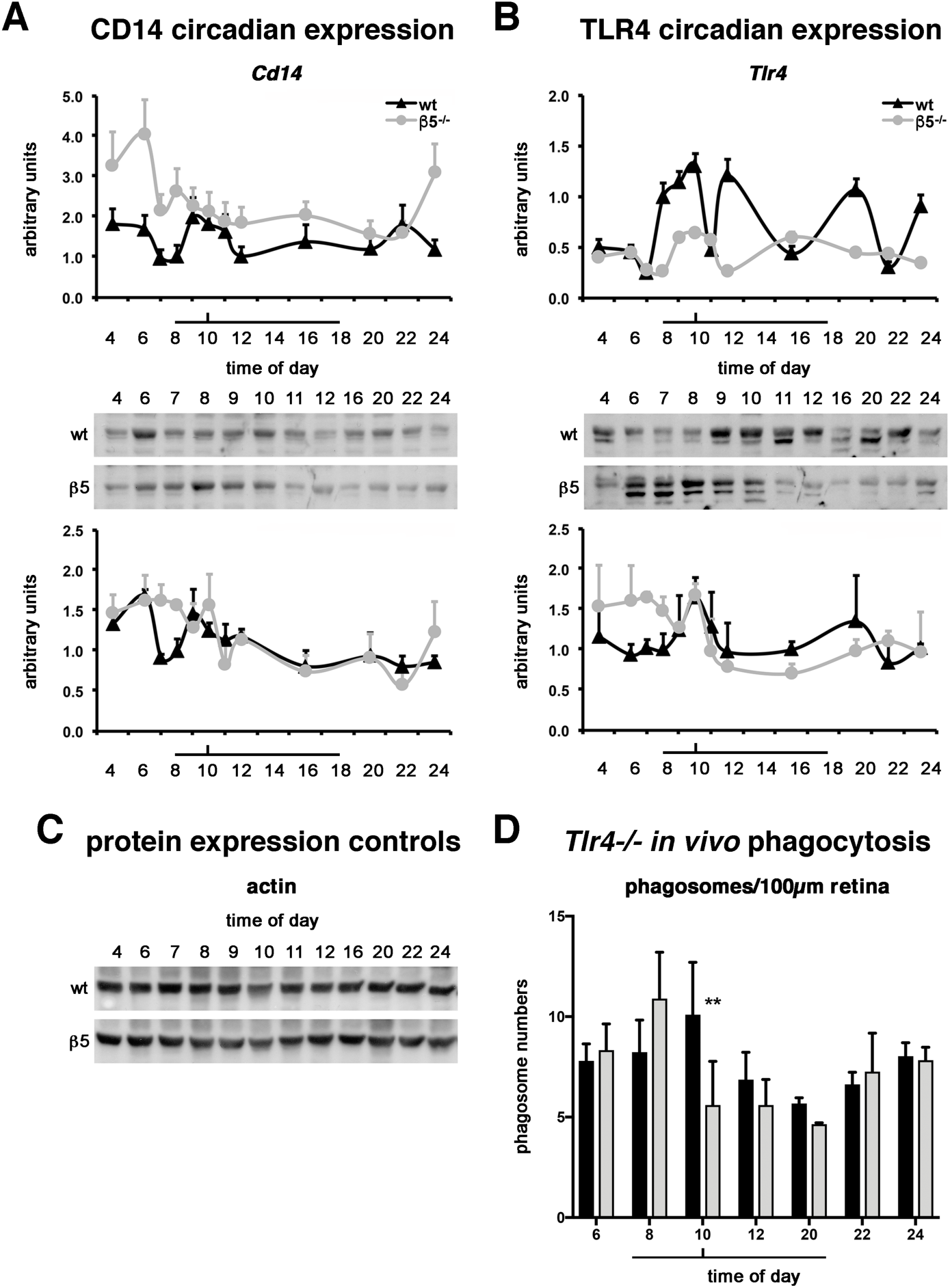
*In vivo* CD14 and TLR4 expression profiles and *Tlr4^-/-^* phagocytic profile. **A,B** CD14 (**A**) and TLR4 (**B**) gene (top) and protein (bottom) expression profiles as shown on graphs and immunoblots in samples from control wildtype (wt, black) and *β5^-/^*^-^ (β5^-/-^, grey) at different times of the light:dark cycle as indicated. RT-qPCR results are expressed in 2^ΔΔCt values. Means ± s.d. in arbitrary units with the reference corresponding to the wt value at 8 hrs (light onset). RT-qPCR *Cd14* N = 7-9, *Tlr4* N = 4-7; immunoblots N = 3-4. **C** Protein expression profiles of actin control in wildtype (wt) and *β5^-/^*^-^ (β5) at different times of the light:dark cycle as indicated. **D** *In vivo* phagocytic profile of wt controls (black bars) and *Tlr4^-/-^*(grey bars) RPE/choroid samples at different times as indicated. The light time period is symbolized by the bar below the graphs, and phagocytic peak time by the vertical tick. Means ± s.d. in phagosome numbers per 100 µm retina, N = 3 (except 20 hrs N = 2), ** P < 0.01 (Two-way ANOVA with Sidak post-correction).

*Tlr4* expression in WT RPE/choroid tissues is upregulated 4 times during the light:dark cycle, between 8 and 10 hrs, at 12 hrs, at 20 hrs and at 24 hrs, while *β5^-/-^* samples show a smooth pattern (Fig. 6B). At the protein level, amounts of TLR4 proteins are augmented at 10 and 20 hrs in WT samples, and show a bimodal pattern in *β5^-/-^* samples with higher expression between 4 and 10 hrs followed by lower expression between 11 and 24hrs. In contrast to profiles observed for CD14 and TLR4 protein levels, protein levels of actin controls do not vary with time in either WT or *β5^-/-^*samples (Fig. 6C).

To fully characterize the role of these receptors *in vivo*, we characterized the phagocytic profile of *Tlr4^-/-^* mice and WT controls at different relevant times of the light:dark cycle on paraffin sections. Interestingly, most of the profile was conserved except at the time of the phagocytic peak that seems to start earlier at 8 hrs and is significantly reduced at the expected peak time at 10 hrs (Fig. 6D).

## Discussion

The molecular machinery mobilized by RPE cells to eliminate daily oxidized POS extremities is tighly controlled as POS and RPE cells are in permanent contact. As studies advance, we have a more and more precise idea of the macromolecular complexes that are involved to keep this process efficient, up and running along the lifespan of the organism. We have already shown the implication of several receptors such as αvβ5 integrin, MerTK and scavenger receptors CD36 and SR-B2/LIMP2 in the regulation and completion of rhythmic POS phagocytosis (Nandrot et al, 2000; Nandrot et al, 2004; Rieu et al, 2022; Enderlin et al, 2024). As RPE cells also expose other phagocytic receptors not explored so far, this work was dedicated to the characterization of the implication of CD14 and TLR4 in POS phagocytosis. It has been shown that CD14 and TLR4 are expressed by the RPE in animal models and human cells (Elner et al, 1981; Elner et al, 2003; Klettner and Roider, 2021). We confirmed their expression in the rat RPE-J cell line and mouse tissues. CD14– and TLR4–POS colocalization during phagocytosis confirms that both receptors recognize POS directly and interact with them. Their surface expression levels increase during phagocytosis suggesting that these receptors are expressed when the cells need them to phagocyte apoptotic POS extremities. *In vivo*, both receptor protein levels increase up to the phagocytic peak or at the time of the peak, confirming this mobilization by the cells to perform phagocytosis.

CD14 and TLR4 are known to work as coreceptors in proinflammatory conditions (Cielseska et al, 2021). We wanted to see if this dynamics takes place in the retina. Our data show that both receptors associate during POS phagocytosis. Furthermore, CD14 and TLR4 colocalization peaks with POS at 3 hrs after POS challenge, suggesting that our receptors may be implicated in mid to late phagocytosis stages, potentially to regulate internalization. Interestingly, we showed that the interaction of these 2 receptors preexists before POS challenge, suggesting the thorough organization of the machinery even before phagocytosis starts.

Inhibition assay data showed that siRNA and protein blockade had different effects when each receptor was blocked alone, targeting CD14 only having an effect and mostly on internalization. CD14 seems to favorise the dynamic of POS phagocytosis since its inhibition, alone or at the same time than TLR4, appears to limit phagocytosis with increased numbers of bound POS concommitant with decreased numbers of internalized POS. This suggests that CD14 might be in more direct contact with POS and its role might be to accelerate POS uptake, an effect also observed for some receptors of the scavenger class A family (Finnemann and Silverstein, 2001; Rieu et al, 2022), but not registered when TLR4 was inhibited alone. However, when using preincubation conditions for TLR4 antibody blockade, internalization was also affected to the same extent than CD14 inhibition, but no cumulative effect was observed. Interestingly, siRNA blocking on J774 macrophages using the same phagocytic machinery had no effect on POS phagocytosis, suggesting a potential tissue-specific regulatory dynamics.

Our cell surface labelings show that both CD14 and TLR4 are organized in clusters at the cell surface, a phenomenon accentuated upon POS challenge. This concentration is certain membrane subdomains suggests that both receptors might be relocating to signaling plateforms such us lipid rafts as was shown for scavenger receptors (Rieu et al, 2022). This concentration in lipid rafts takes place in macrophages upon LPS stimulation (Pleiffer et al, 2001; Płóciennikowska et al, 2015; Ciesielska et al, 2021). Our data on RPE cells show that both receptors colocalize with lipid rafts but with different timings. In fact, it seems that TLR4 is recruited first since the receptor colocalizes with lipid rafts 1 hr after the cells were challenged with POS, when CD14 colocalizes with rafts mostly 3 hrs after POS challenge like class B scavenger receptors do (Rieu et al, 2022). This timing also corresponds to the maximum of CD14-TLR4 colocalization we observe at 3 hrs. In addition, TLR4 seems to stay longer in lipid rafts than CD14, a dynamics correlating with the association and dissociation profile of CD14 and TLR4. These differences suggest that CD14 and TLR4 play different roles in this phagocytic machinery: CD14 seems to be the one recognizing and binding POS first, while TLR4 appears to be more in charge of activating the downstream signaling pathways associated with their interaction. Lipid rafts are considered as signaling platforms facilitating cellular processes by bringing together implicated molecules. In our assays we show that CD14–TLR4 seems to mobilize the MyD88-dependent pathway and not the MyD88-independent TRIF/TRAF pathway. More precisely, the JNK proteins are strongly phosphorylated upon POS challenge, and their activation diminishes with time. In contrast, ERK1/2 (p44/42) proteins activation by phosphorylation increases with time. Hence, these 2 MAPK proteins appear to be sequentially activated, which is interesting as JNK activate TLR-mediated phagocytosis (Ciesielska et al, 2021) and ERK1/2 (p44/42), in association with the PI3K/Akt pathway is involved in cytoskeletal remodeling and phagocytic cup formation in RPE cells (Li et al, 2017), thus both contributing to phagosome internalization. *In vivo*, CD14 and TLR both display a rhythmic expression pattern at the gene and protein levels. The gene expression pattern for *Cd14* is not directly related to the protein expression one, but the *Tlr4* profile is more correlated and RNA expression precedes the proteins synthesis. *Cd14* gene expression peak juste after maximum phagocytosis time, potentially tom reconstitute cellular stoks. CD14 protein expression precedes TLR4, which correlates with our *in vitro* data and the mobilization of CD14 with POS happening before TLR4 is involved. Both receptors are expressed between either before light onset until phagocytosis peak time (1.5-2 hrs after light onset, 10.00AM here), or at peak phagocytosis time. CD14 expression might be regulated by other pathways related to either protein translation by direct synthesis or by export from the internal stores such as the endoplasmic reticulum and the Golgi apparatus, to have the receptors available when required by the cells.

Overall, our study contributed to add other pieces to the puzzle of the complex molecular machinery of retinal phagocytosis. CD14 and TLR4 seem to play a role in the regulation of POS phagocytosis as coreceptors together activating 2 of their associated signaling pathways, but outside of any inflammatory framework in contrast to macrophages. This characteristic reinforces the tissue-specific action in the retina. As well, CD14 and TLR4 act as coreceptors with other regulatory phagocytic receptors such as SR-B2 as shown previously for other scavenger receptors (Amiel et al, 2009), and SR-B2 also interacts with CD36 and the internalization receptor MerTK. We therefore can hypothesize that CD14 and TLR4 might also interact with MerTK. Taken together, the RPE phagocytic machinery in charge of eliminating dozens of POS per day is being more and more revealed as a dynamic macromolecular complex in which multiple molecules intervene, each with a specific role and timing. Interestingly, other receptor associations have been identified in macrophages and involve scavenger receptors, TLR4 (Bowdlish et al, 2009; Amiel et al, 2009; Taban et al, 2022) and CD14 on one hand, and CD14 and MerTK on the other hand (Zizzo and Cohen, 2018). Study of their potentiel interaction in RPE cells may be the key to fully understand the precise sequential activation of this phagocytic machinery and how these different receptors are able to finely regulate this complex function.

## Methods

### Reagents and antibodies

All reagents were purchased from Life Technologies SAS (ThermoFisher Scientific, Courtaboeuf Cédex, France) unless specified. All antibodies and working dilutions used for experiments are detailed in Supplementary Table 1.

### Animals and tissue collection

Integrin beta5 knockout (*β5^-/-^*, RRID:IMSR_JAX:004166) (Huang et al, 2000) and wildtype control (129T2/SvEmsJ; The Jackson Laboratory) mice, as well as Tlr4 knockout (*Tlr4^-/-^*) (Hoshino et al, 1999) and wildtype control (C57BL/6JRj; Janvier Labs) mice were housed under cyclic 12-hour light/12-hour dark conditions (light onset at 8:00) and fed *ad libitum*. Animals were handled following the Association for Research in Vision and Ophthalmology (ARVO) Statement for the Use of Animals in Ophthalmic and Vision Research, and protocols approved by the Charles Darwin Animal Experimentation Ethics Committee from Sorbonne Université and the French Ministry for Higher Education and Research under APAFIS #20191-2019040311402311 v6. For experiments, 3- to 4-month-old mice were euthanized by CO_2_ asphyxiation at different times during the day for the analysis of circadian expression.

The 12 time points analyzed were: 4.00, 6.00, 7.00, 8.00 (light onset), 9.00, 10.00 (rod phagocytosis peak), 11.00, 12.00, 16.00, 20.00 (light offset), 22.00 and 24.00. Eyeballs were carefully enucleated and rinsed in HBSS without Ca^2+^ and Mg^2+^. Retinas were separated from the RPE/choroid layers, and tissues were quickly frozen in liquid nitrogen. For each animal, one eye was used for RNA extraction and gene expression testing, and the fellow eye for protein expression.

For tissue sections, mice were euthanized at 6.00, 8.00, 10.00, 12.00, 20.00, 22.00 and 24.00. Full eyeballs were delicately dissected out and immersed in Davidson fixative for 1 hour at 4°C, a small window in the cornea was made and eyeballs were further fixed in Davidson for 3 hours at 4°C (Enderlin et al, 2024). The anterior segment was dissected out above the iris and samples were placed in Davidson for 3 more hours at 4°C. After overnight dehydration steps using the Spin Tissue Processor STP 120 (Myr, Fisher Scientific SAS, Illkirch, France), eyecups were embedded in paraffin and 5-µm sections deposited on glass slides for further analysis.

### Cell culture

The rat RPE-J cell line (Nabi et al, 1993) was cultivated in DMEM with 4% CELLect Gold FCS (MP Biomedicals, Eschwege, Germany), supplemented with 1 mM Hepes and 1% non-essential amino acids. Cells were maintained in 5% CO_2_ and 32°C. Cells were seeded on coverslips in 24-well plates for immunocytochemistry assays, in 96-well plates for inhibition assays and in 6-well plates for immunoprecipitation assays. Plates were pre-treated with 1% alcian blue 1% acetic acid to enhance adherence. Cells were then polarized for 6 days before being used for experiments. Alternatively, for siRNA inhibition assays cells were cultivated in 96-well plates in DMEM without FCS and allowed to polarize for 4 days.

The J774 mouse macrophage cell line was cultivated in DMEM with 10% FCS supplemented with 1% non-essential amino acids and 1% sodium pyruvate. Cells were maintained in 5% CO_2_ at 37°C.

### POS purification

POS were isolated from the retinas of pig eyes freshly dissected under dim red light as previously described (Parinot et al, 2014). Retinas were collected in a homogenizing solution (20% sucrose, 20 mM tris acetate pH7.2, 2 mM MgCl_2_, 10 mM glucose, 5 mM taurine). The retina suspensions were then thoroughly shaken, filtered and separated onto 25-60% sucrose gradients (tris acetate pH7.2, 10 mM glucose, 5 mM taurine). An ultracentrifugation was then performed at 25,000 rpm for 50 minutes at 4°C. Bright orange bands containing POS were collected and washed in a series of 3 solutions (20 mM tris acetate pH7.2, 5 mM taurine; 10% sucrose, 20 mM tris acetate pH7.2, 5 mM taurine; 10% sucrose, 20 mM Na phosphate pH7.2, 5 mM taurine). Each wash is coupled to a centrifugation step at 5,000 rpm for 10 minutes at 4°C. POS resuspended in DMEM after the last centrifugation were counted, aliquoted and conserved at -80°C with a 2.5% sucrose supplementation. Alternatively, after the resuspension in DMEM POS were labeled using 1 mg/mL fluorescein isothiocyanate (FITC) for 1.5 hours at RT, before proceeding to 2 washes in the third wash solution, DMEM resuspension, counting and aliquoting as described above.

### Phagocytosis assays

Cells were incubated with unlabeled POS resuspended in the appropriate volume of DMEM during 1, 3 and 5 hours for immunocytochemistry and immunoprecipitation assays, and with FITC-labeled POS during 1.5 and 3 hours for phagocytosis quantification of inhibition assays.

For phagocytosis quantification, after POS challenge, cells were washed 3 times with PBS-CM (1X PBS completed with 0.2 mM Ca^2+^, 1 mM Mg^2+^). Half of the cells per plate were then treated with trypan blue in order to quantify internalized POS (Finnemann et al, 1997). Cells were then fixed and nuclei labeled using DAPI. FITC and DAPI numbers were quantified using an automated fluorescent microscope (Arrayscan VTI, HCS Studio software [spot detector v4.1], ThermoFisher Scientific), and FITC/DAPI ratios calculated. Total phagocytosis was quantified from wells not treated with trypan blue, and binding values were obtained by subtracting internalization values from total phagocytosis ratios for each condition (Rieu et al, 2022).

### Sample lysis, immunoprecipitation assays and immunoblotting

Cultured cells and tissues were lysed in 50 mM HEPES pH7.4, 150 mM NaCl, 10% glycerol, 1.5 mM MgCl_2_, 1mM EGTA, 1% sodium deoxycholate, 0.1% SDS and 1% Triton X-100 completed with 1 mM PMSF and 1% each of protease and phosphatase inhibitors (Sigma-Aldrich, Saint Quentin Fallavier, France), phenylmethylsulfonyl fluoride and sodium orthovanadate.

For immunoprecipitation assays, lysates from 6-well plates were collected after phagocytosis and pre-cleared using Protein G agarose beads (Pierce, ThermoFisher Scientific) for 30 min at 4°C on a rotator. After a 5-min spinning step at 7,000 rpm at 4°C, 90% of sample volume were transfered in a new tube and either stored at -80°C. 1-µg of anti–SR-B2 antibody (see Supplementary Table 1) or relevant non-immune IgG was used for the immunoprecipitation step at 4°C for 2 hrs on a rotator. After a 3-min centrifugation at 7,000 rpm at 4°C, Protein G beads were added and allowed to attach precipitates for 1.5 hrs at 4°C on a rotator. 3 washing steps using the lysis buffer without detergents and 3-min centrifugations as detailed above were conducted. Using a pipet, leftover supernatants were removed and reducing sample buffer added to the beads.

Whole cell lysates, isolated cup/retina from one eyecup (12 µg) and immunoprecipitates were separated on 10% SDS-polyacrylamide gels and electroblotted onto nitrocellulose membranes (Whatman, VWR, Rosny-sous-Bois, France). Membranes were probed with primary antibodies overnight at 4°C (see Supplementary Table 1), washed 4 times in 1X TBS 1% Tween-20 at RT and then incubated with secondary antibodies coupled with horseradish peroxidase (1:1,000) for 2 hours at RT. After 4 washes in 1X TBS 1% Tween-20 at RT, chemiluminescence signals on immunoblots were detected using the Western Lightning Plus-ECL reagent (PerkinElmer SAS, Courtaboeuf, France) and chemiluminescence films (Amersham, Dutscher SAS, Bernolsheim, France). After scanning, film scans were treated using the Adobe Photoshop CS6 software, and protein quantification effected using the Image J software (ImageJ, version 1.53t).

### Immunocytochemistry, immunohistochemistry and microscopy

Immunocytochemistry labelings were performed on RPE-J cells after unlabeled POS challenge. Cells were washed 3 times in HBSS-CM (HBSS completed with 0.2 mM Ca^2+^, 1 mM Mg^2+^) on ice. Cell surface proteins were stained using “live labeling” of unfixed cells for 45 minutes on ice with antibodies diluted 1:50 in cold HBSS-CM. Cells were then washed twice in PBS-CM, fixed in 1:10 trichloroacetic acid in ddH_2_O for 15 minutes, and rehydrated for 15 minutes in 30 mM glycine in PBS-CM. Unspecific sites were blocked using 1% BSA in PBS-CM for 30 minutes. Labeling of intracellular proteins was done overnight at 4°C. Cells were then washed twice in 1% BSA in PBS-CM, and stained for 2 hours at RT with appropriate AlexaFluor secondary antibodies (Invitrogen) diluted 1:250. Nuclei were counterstained with DAPI, and coverslips mounting onto glass slides with the FluoroMount-G Mounting Medium (Interchim).

For immuhistochemistry assays, paraffin was removed from eye sections by a 30-minute incubation in SafeSolv solvent substitute (Q Path, VWR, Rosny-sous-Bois, France). Successive rehydrating baths were then performed first for 30 minutes in 100% ethanol, and then 10 minutes each in 90% and 70% ethanol. Unmasking of antibody sites was performed by a 20-minute incubation of the slides in pre-boiled 1X citrate buffer at RT. Pigments were removed using a 5% H_2_O_2_ 1X SSC solution containing deionized formamide under illumination for 20 minutes. After membranes permeabilization with 0.3% Triton in 1X TBS for 5 minutes, non-specific signals were blocked by a 4% BSA solution in 1X TBS. Sections were stained overnight at 4°C with anti-receptor specific antibodies or their respective non-immune IgG controls. Sections were incubated with appropriate AlexaFluor 594 and 647 secondary antibodies diluted 1:1,000 in 1% BSA-TBS for 1 hour at RT, added to a 1:250 dilution of WGA-FITC (Sigma) for rod photoreceptor extracellular matrix staining. Nuclei were counterstained with DAPI, and slides were mounted using Vectashield (Vector Laboratories). Fluorescent images were acquired on an upright Olympus FV1000 confocal microscope equipped with the Fluoview version 2.1c software (Olympus, Rungis, France). Image stacks were treated and compiled using the Fiji (ImageJ, version 2.1.0) and Adobe Photoshop CS6 softwares.

### Inhibition assays

Two types of inhibition assays were performed, siRNA silencing and blocking antibodies. For siRNA inhibition assays, RPE-J cells were double transfected 1 and 3 days post seeding with anti-rat ON-TARGET plus SMARTpool siRNAs (CD14: L-097225-02, TLR4: L-090819-02, Non-targeting control: L-001810-10; Dharmacon), and phagocytosis assays performed on day 4. J774 were transfected 1 day post seeding with anti-mouse ON TARGET plus SMARTpool siRNAs (CD14: L-055665-00, TLR4: L-047487-00; Dharmacon), and tested for phagocytosis on day 4. Efficiency of gene extinction was assessed by RT-qPCR using primers detailed in Supplementary Table 2.

In separate assays, receptors were blocked directly with specific antibodies or their respective non-immune IgG controls at 1 µg/mL during phagocytosis assays, with or without a 1-hour antibody pre-incubation step in DMEM prior to the phagocytic challenge.

### RNA extraction and RT-qPCR

RNAs were extracted from separated retina and RPE/choroid tissues following instructions using 2 DNAse steps as described before (Illustra RNAspin Mini, GE Healthcare, VélizyVillacoublay, France) (Rieu et al, 2022). RNA yields were assessed using a spectrophotometer. 500 ng of RNAs were converted to cDNAs in a 50-µL volume according to the manufacturer’s protocol for 1 hour at 42°C (Reverse Transcription System, Promega, Charbonnières, France). qPCR reactions using the SYBR Green PCR Master Mix were processed on a 7500 Fast Real-Time PCR System apparatus (50°C for 2 minutes, 95°C for 10 minutes, followed by 50 cycles of 95°C for 15 seconds, 60°C for 1 minute) and using the ribosomal protein Rho0 (Rplp0) gene as internal control. Oligonucleotides designed to obtain 150-bp amplicons are listed in Supplementary Table 3. Quantification of target gene expression levels were calculated using the 2^ΔΔCt method, using amounts in wildtype samples at 8 AM (8.00, light onset) as the reference.

### Statistical analysis

All experiments were repeated between 3 and 7 times, except when indicated. The statistical significance of results was determined on the GraphPad prism software using the One-way ANOVA test with a Dunnett post-correction for multiple comparisons (*in vitro*) or the Two-way ANOVA test with a Sidak post-correction for multiple comparisons (*in vivo*), and each row was analyzed individually without assuming a consistent s.d. Significance thresholds of adjusted P values were set as follows: * P < 0.05, ** P < 0.01, *** P < 0.001, **** P < 0.0001.

## Data availability

This study includes no data deposited in external repositories.

## Acknowledgements

The authors would like to thank Océanne Richard, Mathilde Le Maître, Ibtissam Machkour, Dienaba Diakhaby and Abdullah Sham Sher for their contributions to data acquisition for this project during their internships in the laboratory. We also grateful to Anaïs Potey (High-Throughput Screening Facility) and Stéphane Fouquet (Imaging Facility) both at Institut de la Vision for their help with phagocytosis quantification and confocal microscopy, respectively. This work was supported by Agence Nationale de la Recherche [PRC program, ANR-17-CE14-0044-01, Project Grant to E.F.N.], by Laboratory of Excellence Program (LABEX) [LIFESENSES: ANR-10-LABX-65, Project Grant to E.F.N.] and IHU FOReSIGHT [ANR-18-IAHU-0001, Project Grant to E.F.N.] from Agence Nationale de la Recherche, by the Ministère de l’Enseignement Supérieur et de la Recherche [“Contrat Doctoral Handicap” PhD funding to L.D.H.], and by Centre National de la Recherche Scientifique (CNRS) [Tenure Research Director position to E.F.N.]. Additionally, the Institut de la Vision is funded by Institut National de la Santé et de la Recherche Médicale (Inserm), Sorbonne Université and CNRS, and is affiliated to DIM C-BRAINS, funded by the Conseil Régional d’Ile-de-France.

## Author contributions

E.F.N. designed the study; E.F.N., L.D.H., J.E., Q.R., S.K., J.P., C.M. and T.H. performed experiments and collected the data; E.F.N., L.D.H., J.E. and Q.R. analyzed and interpreted the data; G.M. provided the *Tlr4^-/-^* mice; T.H. provided expertise and antibodies validation; E.F.N. and L.D.H. prepared the manuscript. All authors have read and agreed to the published version of the manuscript.

## Supplemental materials

**Supplementary Table 1.**
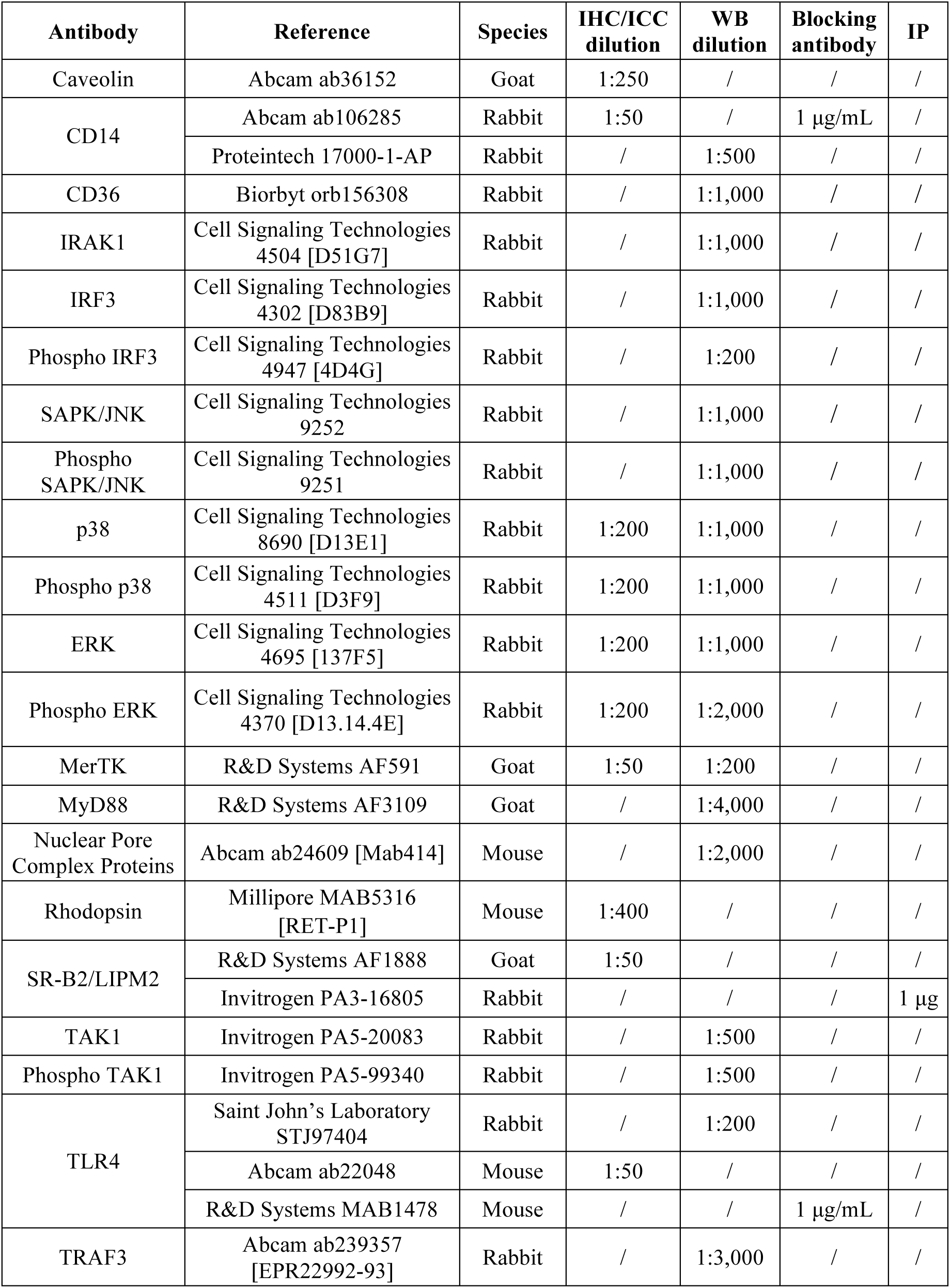
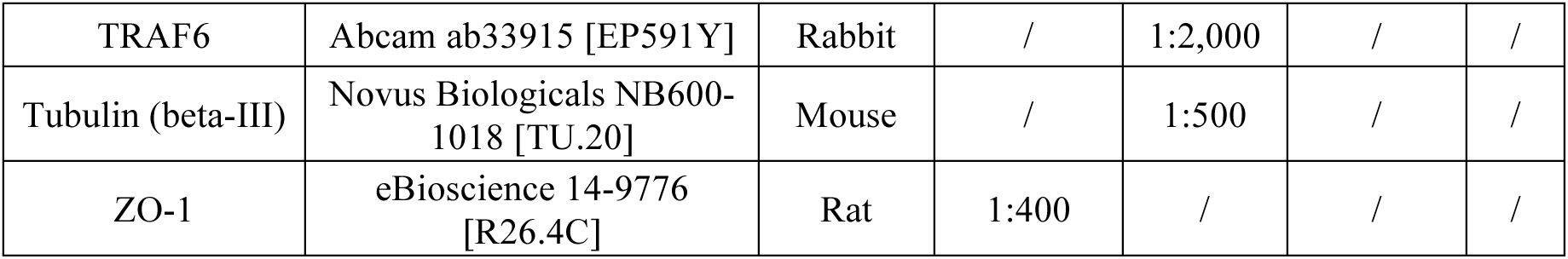
Names, references, species and dilutions corresponding to the antibodies used in immunohistochemical labelings and immunoblots.

**Supplementary Table 2.**
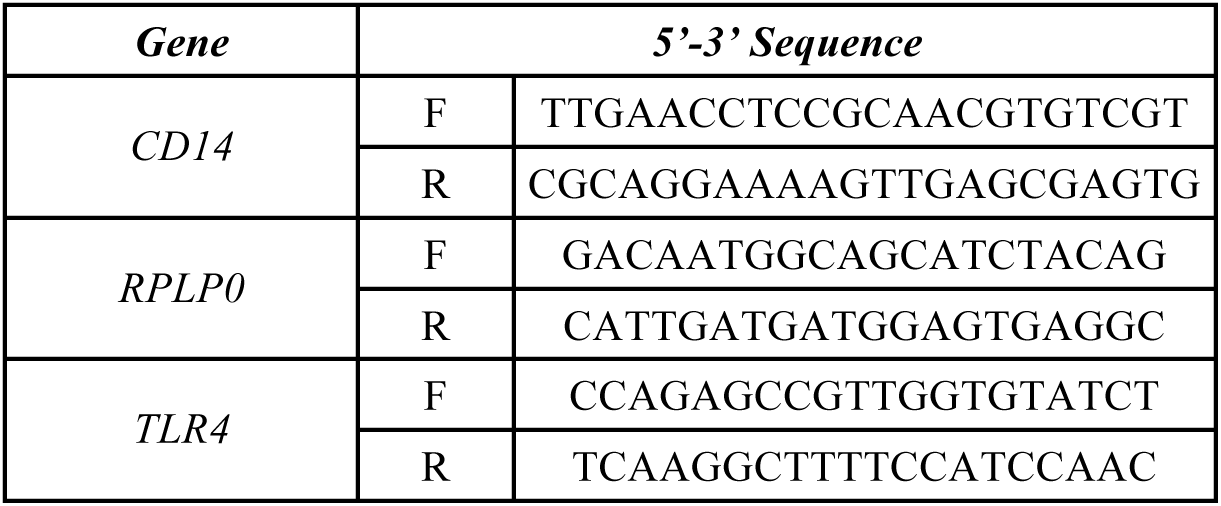
Gene names and corresponding forward (F) and reverse (R) primer sequences used for siRNA validation (rat).

**Supplementary Table 3.**
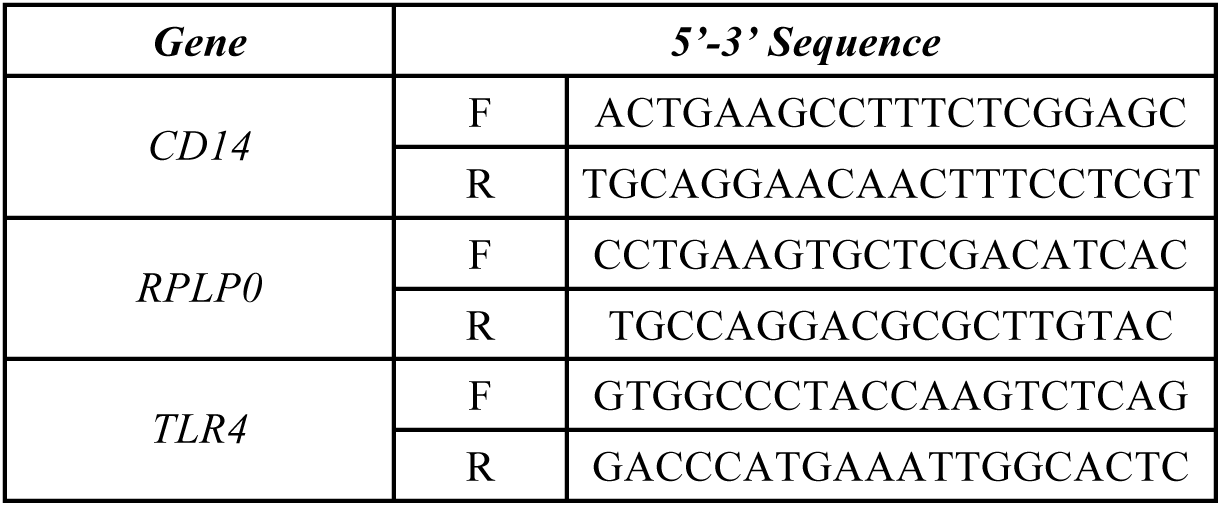
Gene names and corresponding forward (F) and reverse (R) primer sequences used for circadian expression experiments (mouse).

